# The First Insight into the Epigenetic Fusion Gene Landscape of Acute Myeloid Leukemia

**DOI:** 10.1101/2022.12.06.519396

**Authors:** Fei Ling, Noah Zhuo, Degen Zhuo

## Abstract

Epigenetic fusion genes have been defined as the fusion genes produced via *cis*-splicing of read-through pre-mRNAs of two identical-strand neighbor genes and have been known for decades. However, they need to be adequately investigated. In this study, we analyze RNA-Seq data from 390 AML patients and identify 12,754 EFG isoforms encoded by 5,213 EFGs, one-sixth of all potential EFGs. We characterize 479 EFG isoforms whose recurrent frequencies range from 10% to 96.2% and show that most of them result from developmental interactions between recurrent inherited genetic and environmental abnormalities. Novel EFG isoforms generated during late developments reflect somatic genetic abnormalities and environmental stresses. These characteristics of EFG isoforms make it possible for clustering heatmap and counting for EFG isoforms to distinguish GTEx healthy individuals and AML patients. This study reveals that human genomes encode potential EFGs equal to the total number of human genes and pseudogenes. EFGs provide one of the most powerful and economical tools to monitor the earliest signals from somatic genetic and environmental abnormalities.

## Introduction

Acute myeloid leukemia (AML) is a type of blood cancer characterized by excess immature white blood cells^1^. Even though 15-20% of AML cases were diagnosed in children, acute AML incidence was mainly observed in patients over 60 years, and the median age at diagnosis was 70 years old^1^. Environmental factors, including smoking, benzene exposure, and chemotherapy or radiotherapy treatment, increased the risk of developing AML^2,3^. Aging and environmental factors promote somatic genomic abnormalities such as deletions and translocation, resulting in and aggrandizing the cell’s malignant growth^4^. Dissection of developmental interactions between genetics and epigenetics may provide insight into the mechanism of leukemogenesis in AML^3^. Epigenetics has been defined as the study of stable phenotypic changes that do not involve alterations in the DNA sequence^5^. The earlier epigenetic studies focused on regulating access to the DNA template to facilitate transcription, repair, and replication^6^. Read-through fusion genes had been reported in normal cells^7–14^, in cancer cells^7,8,15,16^, and had been systematically annotated for decades^9,17^. However, to discover the cancer-specific fusion genes, many approaches have been developed to remove the fusion genes present in healthy samples from consideration^18–20^, which may hinder our appreciation of them.

We used SCIF (SplicingCodes Identify Fusion Genes) to accurately identify fusion genes, among which *KANSARL* (*KANSL1-ARL17A*) encoded a truncated NSL complex subunit KANSL1 and was validated as the first predisposition fusion gene specific to 29% of European ancestral populations^21^. Validation of highly-recurrent fusion isoforms in healthy samples led us to classify the fusion genes into somatic, inherited, and epigenetic fusion genes^22^. We defined the hereditary fusion genes (HFGs) as the fusion genes offspring inherited from their parents, excluding readthrough^23^. We defined the fusion genes produced via *cis*-spliced fusion isoforms of read-through pre-mRNAs of two identical-strand neighbor genes as epigenetic fusion genes (EFGs)^23^. Our previous study used the curated HFGs uncovered from monozygotic twins^23^ to analyze fusion isoforms and characterized 243 HFGs associated with AML inheritance^22^. In this study, we performed genome-wide analysis and identified one-sixth of potential EFGs, from which 479 EFG isoforms were characterized and ranged from 10% to 96.2% of AML patients. We found that most of these EFGs resulted from developmental interactions between recurrent inherited genetic and environmental abnormalities, and novel EFG isoforms observed during late development were from somatic genetic and environmental lesions. Hence, late EFGs reflected the earliest signals from somatic genetic and environmental alterations during AML initiation and development.

### Identification of epigenetic fusion genes (EFGs) from RNA-Seq of AML patients

Based on their genomic structures, origins, and characteristics, we classified fusion genes into three groups: somatic, hereditary, and epigenetic (read-through) fusion genes^23^. We defined epigenetic fusion genes (EFGs) as *cis*-spliced fusion isoforms of read-through pre-mRNAs of two same-stand neighbor genes. Fig.1a shows examples of EFG isoforms identified in the *NFATC3-PLA2G15-SLC7A6* locus located on 16q22.1. *NFATC3* and *SLC7A6* had been annotated as parts of *DUS2L-NFATC3*, and *SLC7A6-PRMT7* EFGs previously^17^ and here were treated as independent genes to avoid confusion. The most observed EFG types were *NFATC3-PLA2G15^24^* and *PLA2G15-SLC7A6^18^* EFGs (Fig.1a Left and Right), which showed three examples of 22 *NFATC3-PLA2G15* and eight *PLA2G15-SLC7A6* isoforms identified in this study, respectively. Fig.1a (Middle) shows three out of eight *NFATC3-SLC7A6^25^* isoforms, which skipped *PLA2G15* and were a rare EFG type. Fig.1a indicates that the *NFATC3-PLA2G15-SLC7A6* locus generated three EFGs, which encoded enormous isoforms, and suggested that a human genome potentially encoded numbers of EFGs equal human genes. Since gene orders and genomic structures were generally conserved in humans or among different species, our approach was able to identify ≥95% of all potential EFGs. Dissection of all factors inducing EFG expression suggested that EFGs reflected developmentally interactive consequences between environmental stresses and genetic lesions, including inheritance and somatic abnormalities.

**Fig.1.**
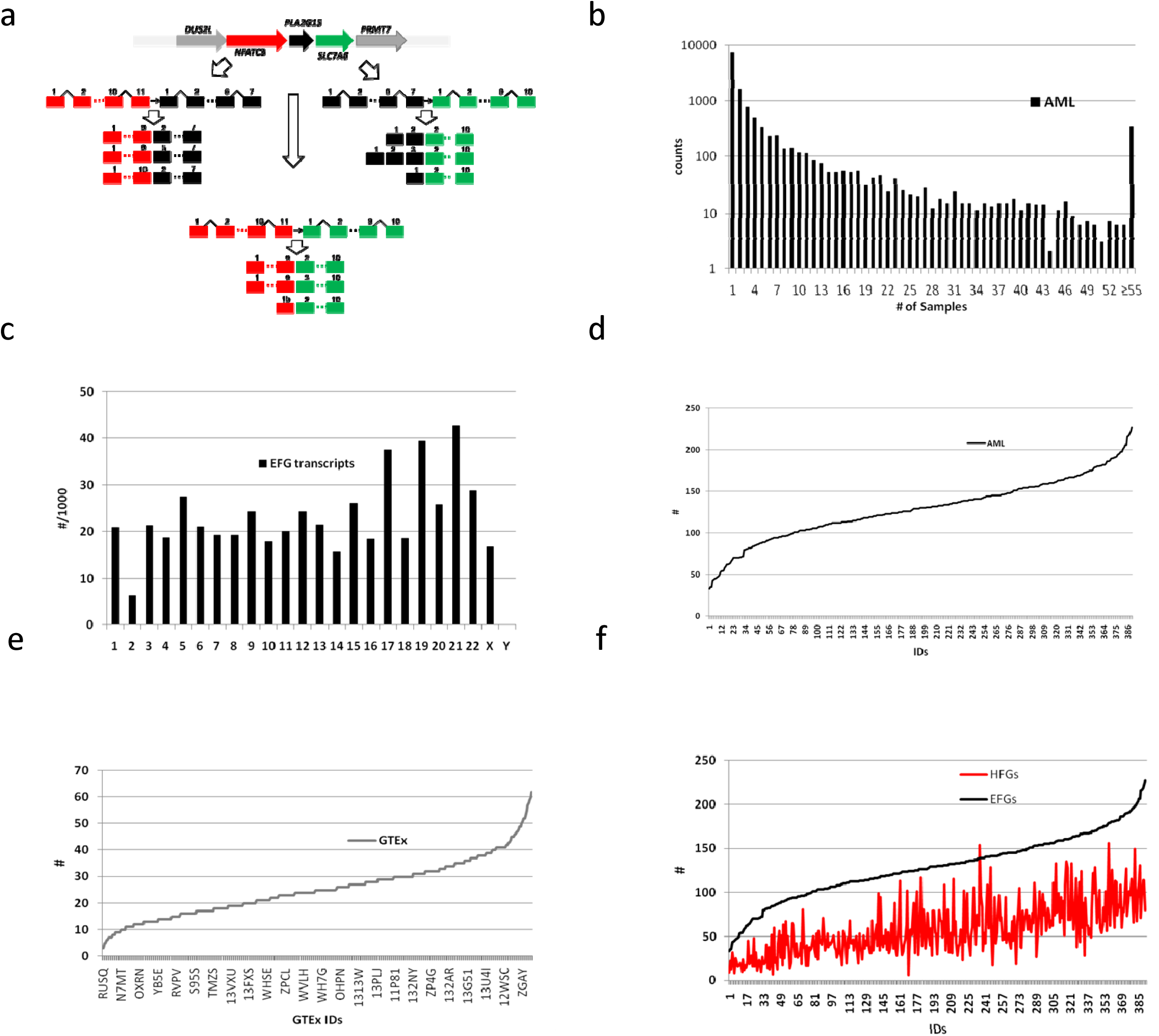
Identification and characterization of Epigenetic fusion genes (EFGs). a). *NFATC3-PLA2G15-SLC7A6* locus was used as an example to demonstrate *cis*-splicing of read-through pre-mRNAs of *NFATC3-PLA2G15-SLC7A6* to produce *NFATC3-PLA2G15, PLA2G15-SLC7A6*, and *NFATC3--SLC7A6* fusion genes. *Red*, black, and green arrows represented *NFATC3, PLA2G15*, and *SLC7A6* genes, respectively. Gray arrows were flanking genes of the *NFATC3-PLA2G15-SLC7A6* locus. Red, black, and green rectangles represented exons of *NFATC3, PLA2G15*, and *SLC7A6* genes, respectively. Angled black lines were intron sequences. Open arrows indicated different types of EFGs; b). The distribution of 12,754 EFG isoforms was coded for by 5,213 EFGs identified from 390 AML patients. The X-axis indicated the number of individuals who possessed a type of EFG isoform. If there was a gap, we added the numbers of EFG isoforms between two X-axis numbers together; c). The distribution of 479 EFG isoforms with recurrent frequencies of ≥10% showed differential expression of EFG isoforms among different chromosomes. The numbers of the X-axis and Y-axis represented human chromosome IDs and numbers of isoforms per 1,000 genes; d). The distribution of numbers of EFG isoforms with recurrent frequencies of ≥10% possessed by 390 AML patients; e). The distribution of numbers of EFG isoforms with recurrent frequencies of ≥10% possessed by 424 GTEx blood samples; f). The relationship between EFG and HFG isoforms with recurrent frequencies of ≥10% possessed by 390 AML patients. The black and red lines represented the EFGs and HFGs.

We used SCIF to analyze RNA-Seq data of 390 AML patients^26^ (GEO: GSE67040) while GTEx’s 424 healthy blood samples (dbGap-accession: phs000424.v7.p2) were used as control, and we identified 1,010,000 fusion isoforms. Systematic examination of all fusion genes showed that distances between two EFG parental genes were most likely within 250,000 bp. Hence, we treated all fusion genes produced from the same stand within 250,000 bp as EFGs. We identified 12,754 EFG isoforms coded for by 5,213 EFGs, which ranged from 1 to 376 of 390 AML patients. Fig.1b shows the frequency distributions of 12,754 EFG isoforms, among which 7191 EFG isoforms were detected only in one AML patient and counted for 56.4% of the total EFG isoforms. Furthermore, 32.7% (1,703) of 5,213 EFGs encoded a single EFG isoform detected only in one AML patient and suggested that they were causal and more likely resulted from random genetic and environmental abnormalities. 43.6% of 12,754 EFG isoforms were observed in ≥2 AML patients and 3.9-fold of the ratio of the total AML fusion isoforms, suggesting that EFGs behaved differently than the other fusion genes. To characterize EFGs more accurately and efficiently, we set the EFG recurrent frequency (RF) cutoff at 10%. Unless they were specified, recurrent frequencies of all EFGs and their isoforms were ≥10% in this study. Hence, we identified 479 EFG isoforms encoded by 374 EFGs, whose recurrent frequencies ranged from 10% to 96.2% (Supplementary Table 1).

To study if EFG expression was associated with chromosomes, we calculated the number of EFG isoforms per 1000 genes for each chromosome. Fig.1c showed that EFG isoforms were plotted against the chromosomes and the numbers of EFGs varied greatly. For example, even though Chromosome 19 had only 163 more genes than Chromosome 2, the EFG isoforms per 1000 genes on Chromosome 19 were 7-fold of their counterparts on Chromosome 2, suggesting that EFGs of Chromosome 19 were activated in AML. These EFG variations among different chromosomes suggested that some EFGs were differentially expressed among the AML chromosomes. To investigate EFG distributions among AML patients, Fig.1d showed that the AML patients had 33 to 227 EFG isoforms, the average of which was 128.9. GTEx blood samples had 16 to 62 EFG isoforms (Fig.1e), the average of which was 25.3. The AML average was five-fold of the GTEx counterpart and suggested that these EFG isoforms were associated with AML. To explore the potential relationship between EFGs and HFGs of AML patients, Fig.1f showed the positive relationship between AML HFG and EFG isoforms, suggesting that the more HFGs AML patients possessed, the more EFG isoforms they had. The average of 129 EFG isoforms per AML patient was 2.2-fold of their HFG counterpart reported previously^22^. The fact that Fig.1f displayed no perfect relationships between EFG and HFG isoforms with recurrent frequencies of ≥10% suggested that other genetic factors and environmental abnormalities were involved in the generation of 479 EFG isoforms.

### Identification and characterization of EFG isoforms associated with AML

To investigate whether individual EFG isoforms were associated with AML, we performed comparative and statistical analyses of 479 EFG isoforms. Supplementary Table 2 showed that nine EFG isoforms were detected in 40% to 80% of GTEx populations and were 40% to 280% higher in GTEx blood samples than the counterparts in AML patients, suggesting that they had a favorable prognosis. The most reduced EFG in AML was *PFKFB4-SHISA5*, which encoded truncated 6-phosphofructo-2-kinase without the last 19 aa C-terminus. *PFKFB4-SHISA5* was detected in 52.9% of GTEx blood samples and 13.8% of AML patients. The former was 2.8-fold of the latter, suggesting that *PFKFB4-SHISA5* had a favorable prognosis for AML. This conclusion was consistent with a previous study showing that blocking 6-phosphofructo-2-kinase in prostate cancer cells allowed free radicals to build up and trigger cell death^27^. The most recurrent EFG among the nine EFGs was *FLJ37307-DHRS12* located on 13q14.3, which coded for a putative transmembrane protein 272-dehydrogenase/reductase 12 (SDS12) hybrid protein and resulted in increased 22 aa at N-terminus of SDS12. The *FLJ37307-DHRS12* isoforms were detected in 80% of GTEx blood samples and were 1.8-folds of the AML counterpart. Since *DHRS12* was one member of >70 human SDS superfamily^28^ and overexpression of *DHRS12* inhibited the proliferation and metastasis of osteosarcoma^29^, *FLJ37307-DHRS12* potentially increased SDS diversities and resulted in tolerance of cellular stresses.

Supplemental Table 3 shows that we identified that 454 EFG isoforms encoded by 354 EFGs were associated with adverse risks to AML patients. They were 30% to 206-fold higher in AML patients than the GTEx counterparts, which were statistically significant. 285 (62.8%) of 454 EFG isoforms ranged from 10-to 206-fold higher in AML than their counterparts in GTEx, suggesting that they were highly recurrent in AML and, at the same time, were rare in GTEx. Supplemental Table 3 showed that *SS18L1-ADRM1* had the highest RF difference between AML and GTEx. *SS18L1-ADRM1* was detected in 48.7% of AML patients and was 206-fold of the GTEx counterpart. *SS18L1-ADRM1* encoded a truncated ADRM1 26S proteasome ubiquitin receptor, shorter than 150 aa and without a cleavable signal peptide and Rpn13 domain-containing protein that was interacted with ubiquitin^30^ and a potential therapeutic target^31^. On the other hand, *PLAUR-CADM4* had the lowest RF difference between AML and GTEx and encoded 618 aa plasminogen activator receptor-cell adhesion molecule 4 hybrid protein. *PLAUR-CADM4* was highly recurrent and detected in 83.8% of AML patients and 63.3% of GTEx individuals, respectively. The former was 32% higher than the latter, suggesting that *PLAUR-CADM4* regulations differed from the *SS18L1-ADRM1*. Supplemental Tables 2&3 showed that the differences between AML and GTEx varied greatly and indicated that the genetic and environmental factors to induce these EFGs were quite divergent and supported large members of HFGs associated with AML^22^.

### Heatmaps of 479 EFG isoforms of AML and GTEx

Next, we used Morpheus to explore 479 EFG isoforms encoded by 374 EFGs. Fig.2a showed that the EFG isoforms were clustered into three groups: the highly-recurrent, moderately-recurrent, and sparsely-recurrent EFG isoforms, which consisted of 10.6%, 24.4%, and 64.9%. The highly recurrent EFG isoforms were from EFG isoforms with recurrent frequencies of ≥54%. The moderately-recurrent isoforms mainly contained EFG isoforms whose recurrent frequencies ranged from 28% to 54%, while the sparsely-recurrent ones were from those with recurrent frequencies of ≤28%. One of the most highly recurrent EFG isoforms was *SLC35A3-HIAT1* encoded fifteen EFG isoforms, six of which were ≥10%. Supplemental Fig.1 showed that *SLC35A3-HIAT1* isoforms a&b resulted in frameshifts and produced different C-termini of solute carrier family 35 member A3. *SLC35A3-HIAT1* isoform *c&d&e* encoded truncated solute carrier family 35 member A3 and were shorter by 82 to 107 aa than solute carrier family 35 member A3. *SLC35A3-HIAT1* isoform *f* encoded solute carrier family 35 member A3-major facilitator superfamily domain containing 14A hybrid protein. Table 1 shows that these six *SLC35A3-HIAT1* isoforms were detected in 14.9% to 73.8% of AML patients while in 0.2% to 3.5% of GTEx blood samples. The formers were 20-to 89-fold higher than the latter, suggesting that *SLC35A3-HIAT1* behaved more like HFGs and had an adverse prognosis to AML. The recurrent frequency differences of *SLC35A3-HIAT1* isoforms supported that *SLC35A3-HIAT1* was a potential pathogenic gene fusion^32^.

**Fig.2.**
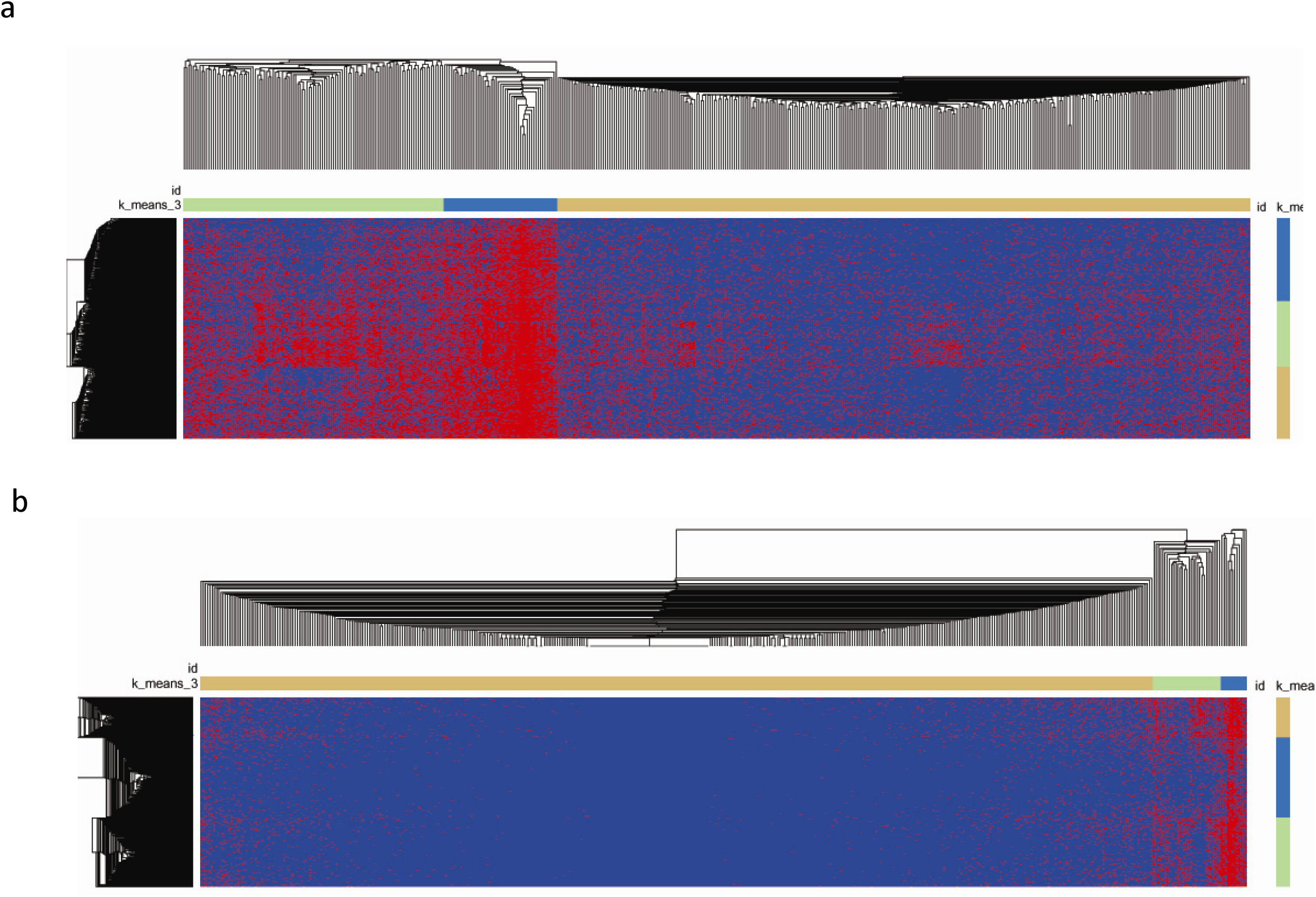
Heatmaps of 479 EFG isoforms generated by Morpheus. a). Heatmap of 479 EFG isoforms with recurrent frequencies of ≥10% among 390 AML patients. The horizontal blue, light-green, and grayorange rectangles represented the highly-recurrent, moderately-recurrent, and sparsely-recurrent EFGs, respectively. The vertical blue, light-green, and gray-orange rectangles showed the AML patients with highly-abundant, moderately-abundant, and sparsely-abundant EFGs, respectively; b). Heatmap of 479 EFG isoforms with recurrent frequencies of ≥10% among 424 GTEx blood samples. The horizontal blue, light-green, and gray-orange rectangles represented the highly-recurrent, moderately-recurrent, and sparsely-recurrent EFGs, respectively. The EFG isoforms possessed by GTEx blood samples were not apparent, as showed by the vertical blue, light-green, and gray-orange rectangles. Horizontal and vertical black lines represented cluster EFG isoforms and AML patient or GTEx healthy blood samples.

**Table 1.**
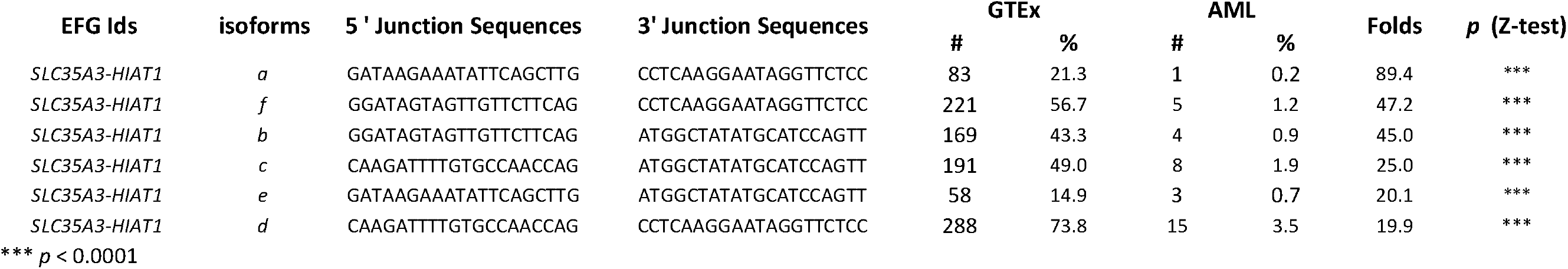
Comparative analysis of SLC35A3-HIAT1 isoforms between AML patients and GTEx samples.

To understand the relationships of AML patients, we used Morpheus to cluster 390 AML patients. Fig.2a showed that 390 AML patients were clustered into three groups: AML patients with the highly-abundant, moderately-abundant, and sparsely-abundant EFGs, which consisted of 29.7%, 33.1%, and 36.2%. These classifications of AML patients based on EFG isoforms were significantly different from their counterparts based on the HFGs, which were 18.5%, 38.2%, and 43.3%, respectively^22^. The increase in the AML patients with the highly-abundant EFG isoforms consisted of a 50% reduction of the AML patients with sparsely-abundant HFGs and a 50% reduction of those with the moderately-abundant HFGs. The AML patients with the highly-abundant EFG isoforms were 60.5% higher than their HFG counterpart^s^^22^, most of which resulted from developmental interactions between recurrent somatic and environmental abnormalities.

To understand overall AML transformation from healthy individuals, we used the identical Morpheus parameters to cluster 479 EFG isoforms from 424 GTEx healthy individuals. As expected, Fig.2b showed that the highly recurrent, moderately recurrent, and sparsely-recurrent EFGs were 2.5%, 6.5%, and 91%, respectively. Most of the sparsely-abundant EFGs were less than 10%, consistent with the above conclusion that 454 EFG isoforms were associated with AML. Fig.2b also showed that when the identical parameters were used to cluster 424 GTEx individuals described above, no groups similar to those of AML patients were observed. The differences between Fig.2a&b suggested that GTEx EFG isoforms reflected the developmentally interactive consequences between genetic and environmental factors. Since GTEx healthy populations were relatively random, their EFGs isoforms were also relatively random. Therefore, the differences in EFG isoforms between GTEx and AML reflected progression from healthy individuals to AML.

### Use of EFG isoforms as biomarkers to diagnose AML

To explore the possible use of 479 EFG isoforms to diagnose AML early, we mixed 390 AML patients with 424 GTEx individuals and then used Morpheus to cluster 814 mixed samples. Fig.3 shows the heatmap generated by Morpheus. Examining detailed clustered data showed that eighteen AML patients were clustered into a subgroup independent from both GTEx and AML groups, indicated by an open arrow in Fig.3 (Supplementary Fig.2), and additional six AML patients were in the GTEx group. On the other hand, only eight GTEx samples overlapped with the fifteen subgroups and the AML group. Hence, Morpheus correctly clustered 94.6% of AML patients and 98.1% of GTEx healthy individuals.

**Fig.3.**
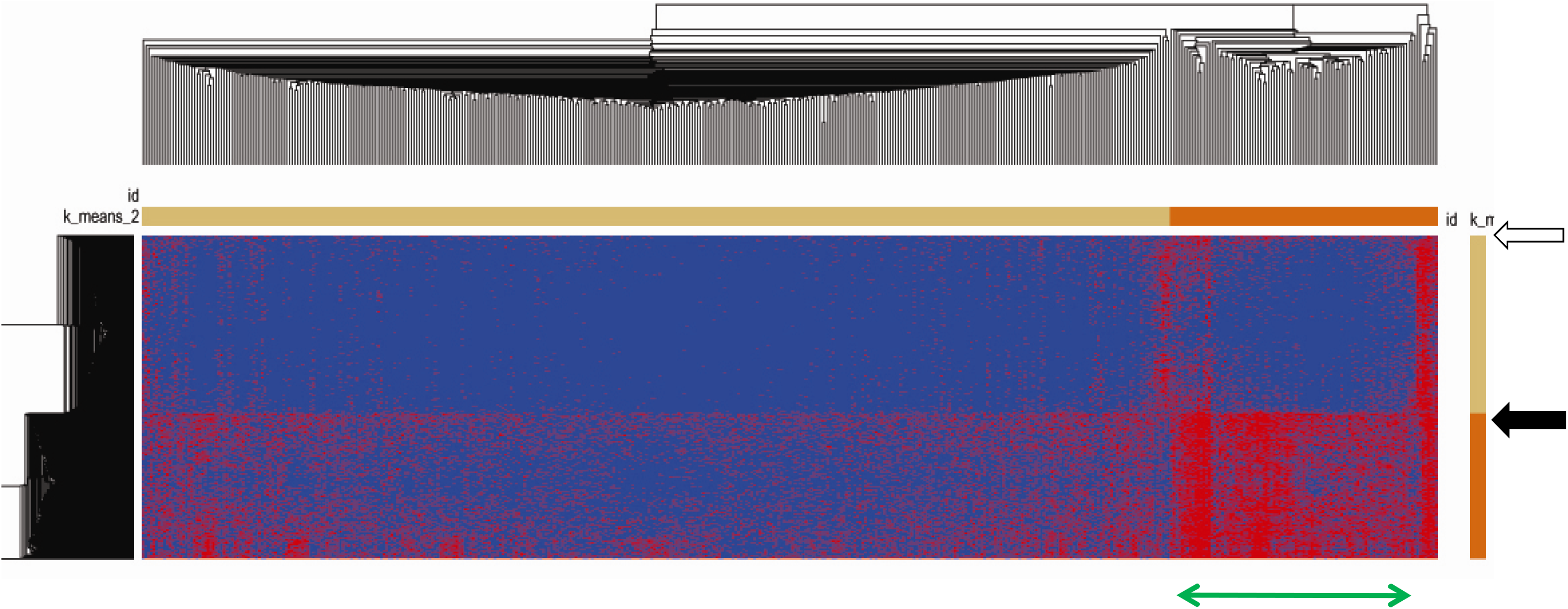
Heatmap of mixed 814 AML patients and GTEx healthy blood samples by Morpheus. The horizontal orange and light-yellow rectangles represented the highly recurrent and sparsely-recurrent EFGs. The vertical orange and light-yellow rectangles showed GTEx healthy blood samples and AML patients. The open arrow indicated a subgroup of AML patients at the top of the heatmap. The black arrow showed the separation between GTEx blood samples and AML patients. Horizontal green arrows indicated the differences between AML patients and GTEx blood samples.

Interestingly, when we directly counted the numbers of 479 EFG isoforms per sample and sorted them based on these raw numbers of EFG isoforms, only nineteen AML patients were mixed in GTEx. Hence, 95.1% of AML patients were correctly predicted based on the 479 EFG isoforms that AML patients had and were even better than the ones clustered by Morpheus. Hence, if we extended other EFG isoforms whose recurrent frequencies were <10%, it would further improve the classification of AML.

## Discussions

Since we defined EFGs as the fusion genes generated via *cis*-splicing of read-through pre-mRNAs of two same-strand neighbor genes (Fig.1a), the maximum numbers of potential EFGs were equal to the number of introns-containing genes, including pseudogenes, which was 35,000^33,34^. Since gene orders and structures were highly conserved, human populations had almost identical numbers of potential EFGs. In this study, 5,213 EFGs were identified and counted for one-sixth of the total potential EFGs, suggesting that most EFGs were not expressed uncontrollably. In contrast to previous reports that EFGs occurred in only 4–6% of the human genome^8,7,9^, our data suggested that nearly all two-same-strand neighbor genes were expressed as EFGs. The machinery regulating EFG expression included transcriptional machinery promoting the expression of 5’ parental genes, components to result in termination failure between two parental genes, and pre-mRNA splicing machinery. All three components had to be correctly aligned simultaneously to express EFGs.

On the other hand, missing any components failed to produce EFGs^35,36^. The factors affecting these three components were developments, environmental stresses, and genetic abnormalities. Based on Fig.1b and EFG data, many EFG isoforms were differentially expressed among different tissues. Hence, one had to be very cautious to make comparison analyses from different tissues. The genetic abnormalities associated with AML consisted of inherited genetic lesions, including HFGs and gene mutations, and somatic genetic alterations composed of somatic fusion genes and gene mutations.

Since HFGs and inherited mutations began interacting with environmental abnormalities from a fertilized egg, EFGs resulting from inherited genetic and environmental lesions appeared earliest and most recurrent. 374 EFGs characterized in this study generally belonged to these EFGs. Since we could use both parents and their offspring to identify HFGs and inherited mutations, these EFG isoforms could discover and study recurrent environmental abnormalities related to AML. During the later AML developmental stages, we identified new sets of total EFGs. The differences in EFG isoforms between the two stages showed reflected developmental interactions between somatic genetic abnormalities and environmental lesions. Fig.2a&b showed the overall increase of highly-recurrent EFG isoforms from healthy individuals to AML and reflected developmental interactions between recurrent genetic and environmental abnormalities. Hence, clustering of EFGs by Morpheus and counting for raw numbers of EFG isoforms in each AML patient had been able to distinguish more than 94.6% of AML patients from healthy individuals. More importantly, novel EFG isoforms between two different stages reflected environmental and somatic genetic abnormalities, which included genomic alterations resulting in the activation of dormant HFGs. In other words, novel EFG isoforms potentially indirectly provided one of the most potent tools to detect somatic gene mutations, genomic abnormalities, and harmful environmental factors.

Fig.1a showed that the *NFATC3-PLA2G15-SLC7A6* locus located on 16q22.1 produced the *NFATC3-PLA2G15 PLA2G15-SLC7A6* and *NFATC3-SLC7A6^25^*, each of which encoded up to 22 isoforms and generated enormous potential diversities of EFG genotypes. Many loci formed synteny and were conserved during animal evolution^37,38^, suggesting that synteny was under functional constraints and prevented random gene reshuffles. One of the most likely constraints was that EFGs from synteny had some essential functions for the viability and inheritability of individuals and new species. For example, the main *CTNNBIP1-CLSTN1* isoform, detected in 61% of AML patients and 5.4% of GTEx blood samples (Supplementary Table 1), had been shown to regulate human neocortical development ^39^. Downregulation of this *CTNNBIP1-CLSTN1* isoform resulted in a marked reduction in human neural progenitors and precocious neuronal differentiation^39^. Fig.2a&b showed that EFG differences between GTEx healthy individuals and AML patients reflected developmental interactions between genetic and environmental abnormalities results. Some EFGs induced by genetic and environmental abnormalities may function as SOS systems^40^ and cell stress responses^41^ to balance harmful effects from genetic and environmental abnormalities. Hence, EFGs deserved to be further explored in the future.

In conclusion, this work was the first attempt to systematically investigate the landscape of AML EFG isoforms. From 12,754 EFG isoforms encoded by 5,213 EFGs, we characterized 479 EFG isoforms with recurrent frequencies ranging from 10% to 96.2%, coded for by 374 EFGs. Detailed analysis showed that these highly-recurrent EFG isoforms were from AML developmental interactions between recurrent inherited genetic factors, most of which were HFGs, and environmental abnormalities. During AML initiation and progression, novel EFG isoforms reflected somatic genetic and environmental abnormalities, providing one of the most potent tools to monitor unknown genetic and environmental abnormalities progressively. These characteristics made it possible for Morpheus and basic counting of EFG isoforms to distinguish GTEx-healthy individuals from AML patients.

## Materials and Methods

### Materials

#### Human acute myeloid leukemia (AML) RNA-Seq dataset

We downloaded RNA-Seq data of the Leucegene AML project^24^ (GEO: GSE67040) from NCBI. The preparation and analysis of RNA-Seq data had been described previously^22^.

#### Genotype-Tissue Expression (GTEx) blood RNA-Seq data

We selected GTEx blood samples as controls to better evaluate the results. We parsed 424 healthy blood samples with unique individual IDs from the GTEx RNA-Seq data (dbGap-accession: phs000424.v7.p2).

### Methods

#### Identification of epigenetic fusion genes

As described previously, we defined epigenetic fusion genes (EFGs) as *cis*-spliced fusion isoforms of read-through pre-mRNAs of two same-stand neighbor genes, whose distances were 250,000 bp^23^. We used SCIF (SplicingCodes Identify Fusion Genes) to analyze RNA-Seq data and identify fusion transcripts. If the two genes of a fusion transcript were located on the identical stand and the distance between the two genes was ≤250,000bp, this fusion transcript was epigenetic.

#### Recurrence frequencies (RFs)

The recurrence frequency (RF) was defined as the number of EFG isoform-positive individuals divided by the total number of samples used in a study.

#### Statistical analysis

We used Z-test to compare recurrent frequency differences between AML patients and GTEx blood samples. If EFG isoforms were detected in AML patients but not in GTEx blood samples, we assumed that GTEx had one sample for calculation convenience. Z-tests were calculated based on the following formula:

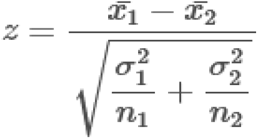

#### Generation of heatmap by Morpheus

Generating heatmaps was performed by Morpheus (https://software.broadinstitute.org/morpheus/) from the Broad Institute. We first used three k-means clustering with Euclidean distance and three members to cluster 479 HFGs and 390 AML patients, and 424 GTEx. Then, we further clustered the EFGs and patients by hierarchical clustering on the K-means. The heatmap was saved as a PDF file.

#### Identification of fusion transcripts by SCIF (<u>S</u>plicing<u>C</u>odes <u>I</u>dentify <u>F</u>usion <u>T</u>ranscripts)

SCIF (<u>S</u>plicing<u>C</u>odes <u>I</u>dentify <u>F</u>usion Transcripts) used to discover fusion genes had been described previously^21^.

## Supporting Materials

Supplementary Table 1: Lists of epigenetic fusion genes (EFGs) with recurrent frequencies of ≥10%

Supplementary Table 2: Lists of epigenetic fusion genes (EFGs) isoforms with favorable AML prognosis

Supplementary Table 3: Lists of epigenetic fusion genes (EFGs) isoforms with adverse AML prognosis

Supplementary Fig.1 Sequence alignments of putative protein sequences from SLC35A3-HIAT1 isoforms. The SLC35A3 protein sequence was used as a reference sequence. Dashed lines indicated alignment sequence gaps.

Supplementary Fig.2. A subgroup independent from both GTEx and AML groups at the heatmap generated by Morpheus. Ids beginning with “SAMN” were AML patients, while the rests of them were GTEx samples.

