## Supplementary Figure 1 for "The First Insight into the Epigenetic Fusion Gene Landscape of Acute Myeloid Leukemia"

Supplementary Fig.1

```
SLC35A3           MFANLKYVSLGILVFQTTSLVLTMRYSRTLKEEGPRYLSSTAVVVAELLKIMACILLVYKD
SLC35A3-HIAT1 a   MFANLKYVSLGILVFQTTSLVLTMRYSRTLKEEGPRYLSSTAVVVAELLKIMACILLVYKD
SLC35A3-HIAT1 b   MFANLKYVSLGILVFQTTSLVLTMRYSRTLKEEGPRYLSSTAVVVAELLKIMACILLVYKD
SLC35A3-HIAT1 c   MFANLKYVSLGILVFQTTSLVLTMRYSRTLKEEGPRYLSSTAVVVAELLKIMACILLVYKD
SLC35A3-HIAT1 d   MFANLKYVSLGILVFQTTSLVLTMRYSRTLKEEGPRYLSSTAVVVAELLKIMACILLVYKD
SLC35A3-HIAT1 e   MFANLKYVSLGILVFQTTSLVLTMRYSRTLKEEGPRYLSSTAVVVAELLKIMACILLVYKD
SLC35A3-HIAT1 f   MFANLKYVSLGILVFQTTSLVLTMRYSRTLKEEGPRYLSSTAVVVAELLKIMACILLVYKD

SLC35A3           SKCSLRALNRVLHDEILNKPMETLKLAIPSGIYTLQNNLLYVALSNLDAATYQVTYQLKIL
SLC35A3-HIAT1 a   SKCSLRALNRVLHDEILNKPMETLKLAIPSGIYTLQNNLLYVALSNLDAATYQVTYQLKIL
SLC35A3-HIAT1 b   SKCSLRALNRVLHDEILNKPMETLKLAIPSGIYTLQNNLLYVALSNLDAATYQVTYQLKIL
SLC35A3-HIAT1 c   SKCSLRALNRVLHDEILNKPMETLKLAIPSGIYTLQNNLLYVALSNLDAATYQVTYQLKIL
SLC35A3-HIAT1 d   SKCSLRALNRVLHDEILNKPMETLKLAIPSGIYTLQNNLLYVALSNLDAATYQVTYQLKIL
SLC35A3-HIAT1 e   SKCSLRALNRVLHDEILNKPMETLKLAIPSGIYTLQNNLLYVALSNLDAATYQVTYQLKIL
SLC35A3-HIAT1 f   SKCSLRALNRVLHDEILNKPMETLKLAIPSGIYTLQNNLLYVALSNLDAATYQVTYQLKIL

SLC35A3           TTALFSVSMLSKKLGVYQWLSLVILMTGVAFVQWPSDSQLDSKELSAGSQFVGLMAVLTAC
SLC35A3-HIAT1 a   TTALFSVSMLSKKLGVYQWLSLVILMTGVAFVQWPSDSQLDSKELSAGSQFVGLMAVLTAC
SLC35A3-HIAT1 b   TTALFSVSMLSKKLGVYQWLSLVILMTGVAFVQWPSDSQLDSKELSAGSQFVGLMAVLTAC
SLC35A3-HIAT1 c   TTALFSVSMLSKKLGVYQWLSLVILMTGVAFVQWPSDSQLDSKELSAGSQFVGLMAVLTAC
SLC35A3-HIAT1 d   TTALFSVSMLSKKLGVYQWLSLVILMTGVAFVQWPSDSQLDSKELSAGSQFVGLMAVLTAC
SLC35A3-HIAT1 e   TTALFSVSMLSKKLGVYQWLSLVILMTGVAFVQWPSDSQLDSKELSAGSQFVGLMAVLTAC
SLC35A3-HIAT1 f   TTALFSVSMLSKKLGVYQWLSLVILMTGVAFVQWPSDSQLDSKELSAGSQFVGLMAVLTAC

SLC35A3           FSSGFAGVYFEKILKETKQSVWIRNIQL-GFFGSIFG--LMGVYIYDGELVSKNGFFQGYN
SLC35A3-HIAT1 a   FSSGFAGVYFEKILKETKQSVWIRNIQL-GFFGSIFG--LMGVYIYDGELVSKNGFFQGYN
SLC35A3-HIAT1 b   FSSGFAGVYFEKILKETKQSVWIRNIQL-GFFGSIFG--LMGVYIYDGELVSKNGFFQGYN
SLC35A3-HIAT1 c   FSSGFAGVYFEKILKETKQSVWIRNIQLDGYMHPVHPHVVRRLAYQHNQTVCRNSPTVEM-
SLC35A3-HIAT1 d   FSSGFAGVYFEKILKETKQSVWIRNIQL-GFFGSIFG--LMGVYIYDGELVSKNGFFQGYN
SLC35A3-HIAT1 e   FSSGFAGVYFEKILKETKQSVWIRNIQL----------------------ASRNRFS----
SLC35A3-HIAT1 f   FSSGFAGVYFEKILKETKQSVWIRNIQL-GFFGSIFG--LMGVYIYDGELVSKNGFFQGYN

SLC35A3           RLTWIVVVLQALG----GLVIAAVIKYADNILKGFATSLSIILSTLISYFWLQDFVPTSV-
SLC35A3-HIAT1 a   RLTWIVVVLQALG----GLVIAAVIKYADNILKGFATSLSIILSTLISYFWLQDFVPTSLK
SLC35A3-HIAT1 b   RLTWIVVVLQALG----GLVIAAVIKYADNILKGFATSLSIILSTLISYFWLQDFVPT---
SLC35A3-HIAT1 c   -------------------------------------------------------------
SLC35A3-HIAT1 d   RLTWIVVVLQ--M----AICIQFILM-----------------------------------
SLC35A3-HIAT1 e   -------------------------------------------------------------
SLC35A3-HIAT1 f   RLTWIVVVLQPQGIGSPSVYHAVIVIFLEFFAWGLLTAPTLVV--LHETFPKHTFLMNGLI

SLC35A3           -FFLGAILVITATF----LYGYDPKPAGNPTKA----------------------------
SLC35A3-HIAT1 a   EVLLVSIMQLSSSFWSFLLGDY---------------------------------------
SLC35A3-HIAT1 b   RWLYAS----SSSSCS---------------------------------------------
SLC35A3-HIAT1 c   -------------------------------------------------------------
SLC35A3-HIAT1 d   -------------------------------------------------------------
SLC35A3-HIAT1 e   -------------------------------------------------------------
```

```
SLC35A3-HIAT1 f  QGVKGLLSFLSAPLIGALSDVWGRKSFLLLTVFFTCAPIPLMKISPWWYFAVISVSGVFAV

SLC35A3          -------------------------------------------------------------
SLC35A3-HIAT1 a  -------------------------------------------------------------
SLC35A3-HIAT1 b  -------------------------------------------------------------
SLC35A3-HIAT1 c  -------------------------------------------------------------
SLC35A3-HIAT1 d  -------------------------------------------------------------
SLC35A3-HIAT1 e  -------------------------------------------------------------
SLC35A3-HIAT1 f  TFSVVFAYVADITQEHERSMAYGLVSATFAASLVTSPAIGAYLGRVYGDSLVVVLATAIAL

SLC35A3          -------------------------------------------------------------
SLC35A3-HIAT1 a  -------------------------------------------------------------
SLC35A3-HIAT1 b  -------------------------------------------------------------
SLC35A3-HIAT1 c  -------------------------------------------------------------
SLC35A3-HIAT1 d  -------------------------------------------------------------
SLC35A3-HIAT1 e  -------------------------------------------------------------
SLC35A3-HIAT1 f  LDICFILVAVPESLPEKMRPASWGAPISWEQADPFASLKKVGQDSIVLLICITVFLSYLPE

SLC35A3          -------------------------------------------------------------
SLC35A3-HIAT1 a  -------------------------------------------------------------
SLC35A3-HIAT1 b  -------------------------------------------------------------
SLC35A3-HIAT1 c  -------------------------------------------------------------
SLC35A3-HIAT1 d  -------------------------------------------------------------
SLC35A3-HIAT1 e  -------------------------------------------------------------
SLC35A3-HIAT1 f  AGQYSSFFLYLRQIMKFSPESVAAFIAVLGILSIIAQTIVLSLLMRSIGNKNTILLGLGFQ

SLC35A3          -------------------------------------------------------------
SLC35A3-HIAT1 a  -------------------------------------------------------------
SLC35A3-HIAT1 b  -------------------------------------------------------------
SLC35A3-HIAT1 c  -------------------------------------------------------------
SLC35A3-HIAT1 d  -------------------------------------------------------------
SLC35A3-HIAT1 e  -------------------------------------------------------------
SLC35A3-HIAT1 f  ILQLAWYGFGSEPWMMWAAGAVAAMSSITFPAVSALVSRTADADQQGVVQGMITGIRGLCN

SLC35A3          -------------------------------------------------------------
SLC35A3-HIAT1 a  -------------------------------------------------------------
SLC35A3-HIAT1 b  -------------------------------------------------------------
SLC35A3-HIAT1 c  -------------------------------------------------------------
SLC35A3-HIAT1 d  -------------------------------------------------------------
SLC35A3-HIAT1 e  -------------------------------------------------------------
SLC35A3-HIAT1 f  GLGPALYGFIFYIFHVELKELPITGTDLGTNTSPQHHFEQNSIIPGPPFLFGACSVLLALL

SLC35A3          -----------------------------------------------
SLC35A3-HIAT1 a  -----------------------------------------------
SLC35A3-HIAT1 b  -----------------------------------------------
SLC35A3-HIAT1 c  -----------------------------------------------
SLC35A3-HIAT1 d  -----------------------------------------------
SLC35A3-HIAT1 e  -----------------------------------------------
SLC35A3-HIAT1 f  VALFIPEHTNLSLRSSSWRKHCGSHSHPHNTQAPGEAKEPLLQDTNV
```

Supplementary Fig.1  Sequence alignments of concepted protein sequences from SLC35A3-HIAT1 isoforms.
The SLC35A3 protein sequence was used as a reference sequence. Dashed lines indicated alignment sequence
gaps.
