## Supplementary Figure 2 for "The First Insight into the Epigenetic Fusion Gene Landscape of Acute Myeloid Leukemia"

S Fig.2

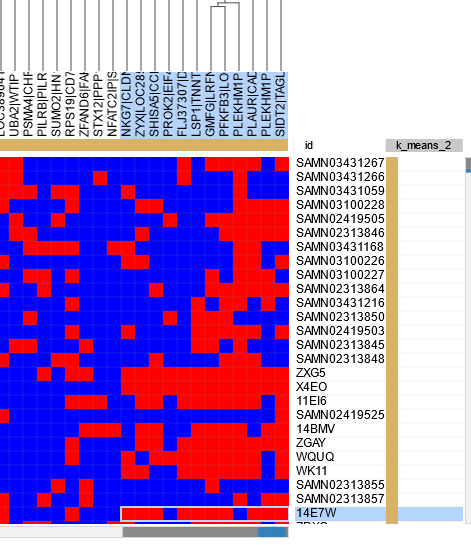

Supplementary Fig.2. A subgroup independent from both GTEx and AML groups at the heatmap generated by Morpheus. Ids beginning with “SAMN” were AML patients while the rests of them were GTEx samples.
